# Marketplace Shrimp Mislabeling in North Carolina

**DOI:** 10.1101/734376

**Authors:** Morgan L. Korzik, Hannah M. Austin, Brittany Cooper, Caroline Jasperse, Grace Tan, Emilie Richards, Erin T. Spencer, Blaire Steinwand, F. Joel Fodrie, John F. Bruno

## Abstract

Seafood mislabeling occurs in a wide range of seafood products worldwide, resulting in public distrust, economic fraud, and health risks for consumers. We quantified the extent of shrimp mislabeling in coastal and inland North Carolina. We used standard DNA barcoding procedures to determine the species identity of 106 shrimp sold by 60 vendors across North Carolina as “local” shrimp. Thirty-four percent of the purchased shrimp was mislabeled, and surprisingly the percentage did not differ significantly between coastal and inland counties. Roughly one third of product incorrectly marketed as “local” was in fact whiteleg shrimp: an imported, and very likely farmed, species from the eastern Pacific (and not found in North Carolina waters). In addition to the negative ecosystem consequences of shrimp farming (e.g., the loss of mangroves forests and the coastal buffering they provide) and seafood importation, North Carolina fishers—as with local fishers elsewhere—are negatively impacted when vendors label farmed, frozen, and imported shrimp as local, fresh, and wild-caught.

## Introduction

Shrimp is the most popular seafood in the United States. Shrimp represents a quarter of America’s annual per capita seafood consumption and the average American eats about four pounds of shrimp a year (NMFS, 2015). This results in over one billion pounds of shrimp consumed annually in the United States alone (NMFS, 2015). Despite a robust domestic shrimp fishery, 90% of the shrimp consumed in the U.S. is imported (NMFS, 2015). In 2018, the U.S. imported 68,000 tons of shrimp, primarily from Indonesia, India, and Ecuador, which accounted for 33% of all seafood imports (NMFS, 2019; NMFS, 2015).

Shrimping has deep cultural and economic roots on the North Carolina coast. In 2017, commercial fishermen caught 13.9 million pounds of shrimp, which is 82.9% greater landings than the previous five year average (N.C. DMF, 2018). That same year, shrimp was the highest earning fishery in the state, valuing $29.6 million, and were exceeded in catch weight only by blue crab (N.C. DMF, 2018). Despite a 9% annual decline in commercial landings for all species in North Carolina in 2017, the value of local shrimp was at an all-time high, suggesting sustained demand. However, a growing threat to North Carolina’s seafood industry is imported seafood products. The number of licensed commercial fishermen in North Carolina declined 41% between 1995 and 2011, and the number of seafood processors on the eastern shore of the state declined 36% between 2000 and 2011 (Kros and Rowe, 2013; Garrity-Blake and Nash, 2012). Despite the increasing prevalence of imported seafood, 92% of North Carolina consumers surveyed by NC Sea Grant indicated they prefer to eat local seafood over imported seafood (Nash and Andreatta, 2011).

In response to increasing pressure on commercial fisheries from seafood imports, organizations in North Carolina and other states have conducted outreach to consumers encouraging them to “eat local seafood”. For example, NC Catch is a local seafood advocacy organization in North Carolina that provides information on vendors that sell local seafood. The intent is to give consumers the tools to make informed decisions regarding seafood purchases that support the local fishing industry. However, this is only effective when seafood products are represented accurately.

Seafood mislabeling occurs when a species is substituted with another type of seafood, including ones of lower economic value (Marko et. al, 2004). Commercial catch restrictions on in-demand species can create an economic incentive for vendors to sell lower-valued fish as more expensive ones (Marko et. al, 2004). Seafood mislabeling can occur at any point along the supply chain, from initial harvest to consumer purchase. It can be difficult to determine where in the supply chain mislabeling has occurred, allowing the practice to continue despite growing public awareness (Cawthorn et al., 2018).

According to a 2014 study, between 30% and 38% of seafood is mislabeled in the United States, and 30% is mislabeled worldwide (Warner et al., 2014; Warner et al., 2016). Mislabeling has a myriad of potential consequences, including exacerbating over-fishing, negative impacts on human health, and perpetuating human rights abuses in international fisheries (Cox et al., 2013; Marko et al., 2014; Kittinger et al., 2017). Understanding more about the scope and frequency of mislabeling can help pinpoint the sources and more effectively monitor and enforce mislabeling practices. As seafood products can be difficult to distinguish visually, an increasing number of studies are using DNA barcoding to quantify the frequency of mislabeling across different species and geographic regions (Willette et. al, 2017).

We used standard DNA barcoding techniques to quantify the level of shrimp mislabeling in North Carolina. Few studies have focused specifically on shrimp mislabeling, and none have assessed the prevalence of shrimp mislabeling in North Carolina. We considered three shrimp species, *Farfantepenaeus aztecus, Litopenaeus setiferus*, and *Farfantepenaeus duorarum*, as “local” to North Carolina. *F. aztecus*, or brown shrimp, is the most abundant shrimp species in North Carolina and accounts for 67% of the state’s shrimp catch (N.C. Division of Marine Fisheries). *L. setiferus*, or white shrimp, is the second-most abundant shrimp species and accounts for approximately 28% of shrimp landings (N.C. Division of Marine Fisheries). *F. duorarum*, or pink shrimp, only account for 5% of the state’s shrimp catch (N.C. Division of Marine Fisheries).

## Materials and Methods

To determine the frequency of shrimp mislabeling in North Carolina, we collected “local” shrimp sold at coastal and inland vendors, including grocery stores and seafood-specific markets. All vendors reported to NC Catch that they sold seafood caught in North Carolina. Some shrimp had signage that explicitly labeled them as local, while others were verified as local by the retail personnel. Samples were only collected when the vendor explicitly or verbally confirmed the product was local or North Carolina shrimp. We obtained samples from 60 vendors across North Carolina (31 were inland vendors and 29 were coastal) in summer and fall of 2017 and during the summer of 2018 (Fig. 1). We defined inland vendors as ones located in a land-locked county. Of 106 total samples processed, 47 were from inland vendors, and 59 were from coastal vendors. Most locations were only sampled once, yet multiple samples were taken from vendors that sold various types of shrimp, including veined/deveined and small/medium/large. Additionally, ten vendors were revisited from 2017 to 2018 to check for mislabeling consistency. We obtained three separate shrimp from each sample and froze these in 2-mL scintillation vials for later storage. We only performed DNA extraction on the first vial unless failure in DNA extraction or amplification necessitated extraction from another sample. When purchasing shrimp we also recorded the cost ($/lb) for each sample collected.

**Fig. 1.**
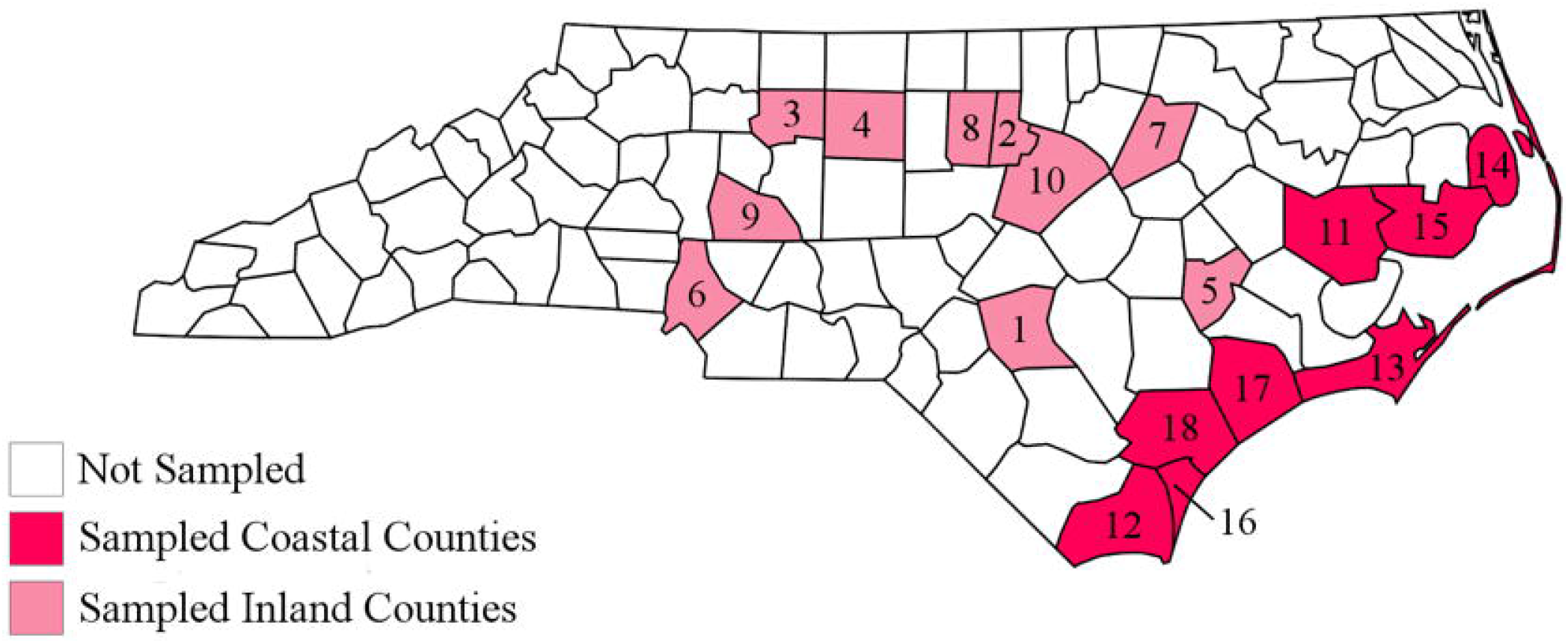
Distribution of sampled vendors in North Carolina. The inland counties include Cumberland (1), Durham (2), Forsyth (3), Guilford (4), Lenoir (5), Mecklenburg (6), Nash (7), Orange (8), Rowan (9), and Wake (10). The coastal counties include Beaufort (11), Brunswick (12), Carteret (13), Dare (14), Hyde (15), New Hanover (16), Onslow (17), and Pender (18).

Following the DNAEasy extraction protocol (Quiagen, INC), we extracted genomic DNA from approximately 20mg shrimp tissue. To identify individual samples to the species-level we focused on sequencing the mitochondrial DNA cytochrome oxidase I gene (CO1). This gene is well-conserved, has little variation within a species, and has enough variation between species to make it a good candidate for our study. It has been used in other seafood mislabeling studies (e.g., Cox et. al 2013, Staffen et al. 2017, Willette et. al 2017). We amplified CO1 sequences from extracted DNA following the PCR protocol outlined in Willette et al. (2017) and a primer cocktail from Ivonova et al. (2008). To prepare a 25μL sample for PCR, we combined the DNA with a primer cocktail of CO1_F1, CO1_F2, CO1_R1, and CO1_R2, deionized water, and a PuRe Taq Ready-To-Go PCR bead containing the necessary PCR components. In the thermal cycler, the samples went under 35 cycles of 95°C for denaturation, 50°C for annealing, and 70°C for extension. A negative control containing all of the PCR components, except DNA, was used to test for contamination. We ran the PCR products on a 1% agarose gel to determine if PCR amplification of the DNA was successful. Samples with successful ~650 base pair bands were sent to an ETON Bioscience facility in Raleigh, NC for sequencing. Chromatograms of successfully sequenced regions were then matched against CO1 sequences of known samples on National Center for Biotechnology Information’s nucleotide collection database GenBank using the Basic Local Alignment Search Tool (BLAST). We only concluded the identity of a species if the percent identity and query coverage was greater than or equal to 98% and the e-value was close to zero. Samples identified as *Litopenaeus setiferus* were considered to be local North Carolina shrimp while *Litopenaeus vannamei* samples were determined to be Pacific whiteleg shrimp, and thus mislabeled.

## Results

Of 128 total samples collected, 106 samples were successfully sequenced. Thirty six of the 106 processed shrimp (34%) were identified as *Litopenaeus vannamei* and 70 were *Litopenaeus setiferus*. These were the only two shrimp species identified in our study. *L. setiferus* or “white shrimp” are native to the western Atlantic and are harvested along the eastern coast of the United States, and in the Gulf of Mexico. *L. vannamei*, whiteleg shrimp, is native to the eastern Pacific. It is common farmed, especially in coastal Ecuador (Fofonoff et. al, 2018). Shrimp sold as “local” and identified as *L. vannamei* were considered mislabeled.

There was no statistical difference in mislabeling frequency between coastal and inland vendors (χ^2^=0.212 and p=0.65). 35% of vendors mislabeled local shrimp at least once. Of ten resampled vendors, six sold both correctly labeled and mislabeled shrimp. The price of mislabeled shrimp (mean: $11.00/lb) was significantly lower than that of the correctly labeled samples (mean: $13.20/lb, p-value=0.001, t-test, Fig. 2).

**Fig. 2.**
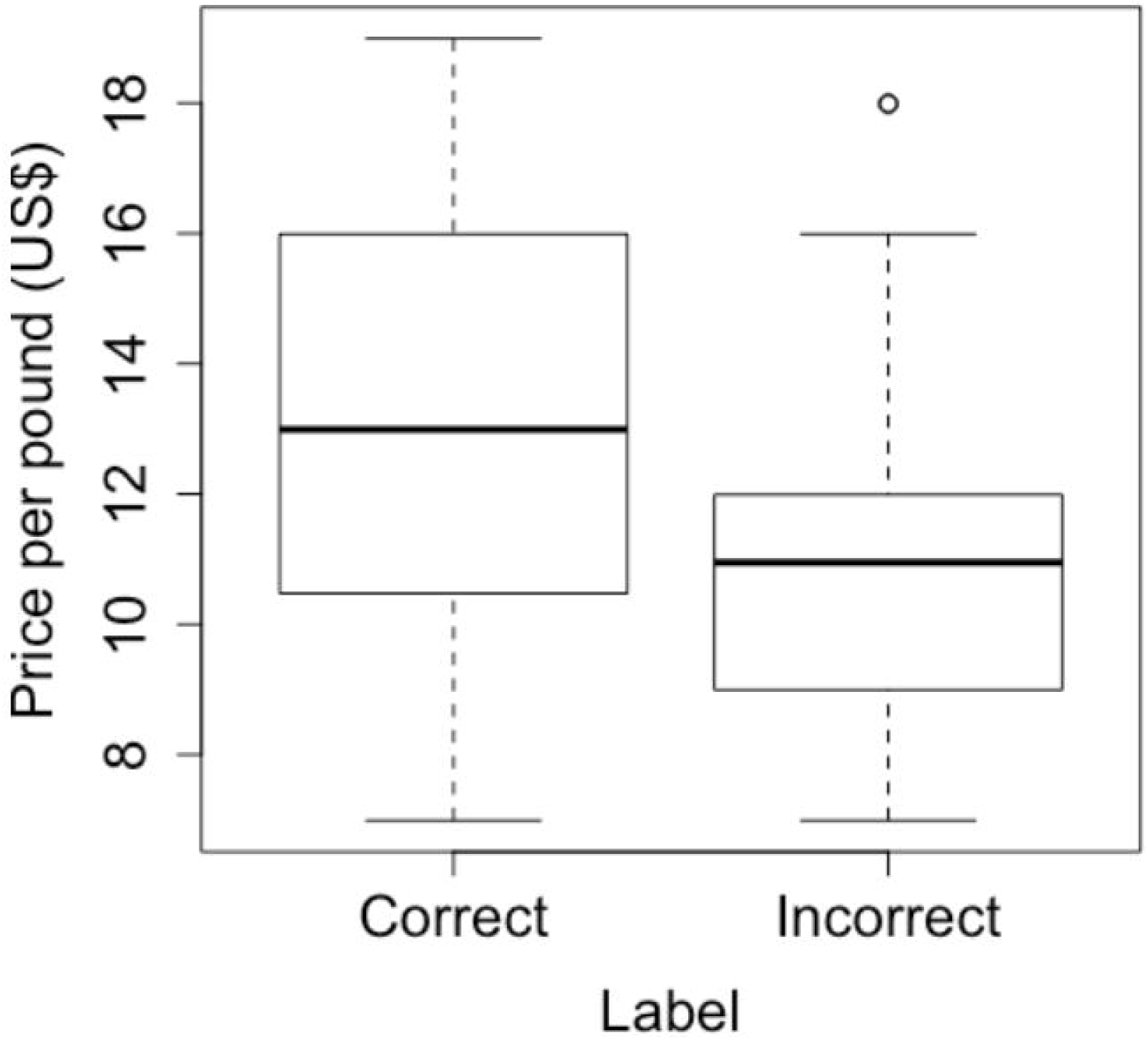
Price of correctly labeled and mislabeled shrimp purchased in North Carolina. Boxplots of the distribution of price per pound of all shrimp that were found to be correctly labeled as “local” compared the price of shrimp incorrectly labeled as “local”. The difference was statistically significant (p-value=0.001, t-test).

## Discussion

Of the 60 sampled vendors across North Carolina, 35% mislabeled local shrimp at least once. This statewide mislabeling frequency is consistent with the 35% shrimp mislabeling frequency nationwide (Warner et al., 2014). Although this frequency is lower than that of other species in North Carolina, e.g., red snapper, (Spencer and Bruno, in review) the results suggest shrimp mislabeling is a fairly common problem across the state.

Shrimp mislabeling may have both ecological, economical, and human health impacts. Locally-caught wild white shrimp are considered a smart seafood choice by the National Oceanic and Atmospheric Association (NOAA) because it is sustainably harvested and managed in the South Atlantic and Gulf of Mexico (NOAA, 2018). White shrimp populations are above target levels, and gear restrictions, such as the required inclusion of turtle excluder devices and bycatch reduction devices, are in place to minimize impacts of trawling on benthic ecosystems (NOAA, 2018).

Pacific whiteleg shrimp are harvested through trawling, which can be destructive to benthic ecosystems and result in high levels of bycatch without gear restrictions like those present in the United States (Clucas, 1997; Lobo et. al, 2010). Additionally, Pacific whiteleg shrimp is the most widely farmed shrimp species in the world and is cultivated in at least 27 countries (Fofonoff et. al, 2018). Shrimp farming poses a number of environmental risks, including mangrove destruction and the associated loss of native biodiversity and ecosystem services (Paul and Vogl, 2011). Shrimp farms often use large doses of antibiotics to prevent the spread of disease, which can contribute to antibiotic resistance in both shimp and human populations. Some antibiotics used, such as enrofloxacin and chloramphenicol, are not advised for human use due to risks of cancer and immune system damage (Avila, 2012). A study in Thailand found 74% of interviewed shrimp farmers used up to 13 different antibiotics in their shrimp ponds, sometimes daily, and many were poorly informed about safe application of antibiotics (Holmstrom et al., 2003).

There are also human rights concerns with imported shrimp (Kittinger et al. 2017). In 2014, multiple news organizations reported slavery practices on Thai fishing vessels harvesting offshore fish to use for farmed shrimp feed (Marschke and Vandergeest, 2016). Exposure of these practices led to a consumer movement to eat seafood that was both environmentally sustainable and ethically harvested. Mislabeling imported shrimp as locally-caught shrimp undermines the power of the consumer to spend their money in a way that aligns with their moral principles. This also fosters consumer distrust in the seafood industry, which could lead to decreased spending on seafood products.

Surprisingly there was no difference in mislabeling frequency between coastal and inland vendors. Of the 10 vendors sampled in both 2017 and 2018, six sold both correctly-labeled and mislabeled shrimp across different years. Further research is needed to know whether there are temporal trends in mislabeling: revisiting vendors throughout the year could help determine whether mislabeling frequency changed based on seasonal fishery closures or tourism activity. Mislabeling frequency could be higher when market demand is high, yet the commercial shrimp fishery is closed or when the shrimp are out of season.

## Acknowledgements

This study was supported by the Department of Biology at The University of North Carolina at Chapel Hill and funded by the QEP (Quality Enhancement Plan) CURE (Course-based Undergraduate Research Experience) initiative. Samples were collected and analyzed as part of the Seafood Forensics CURE class at UNC-CH, specifically students in summer and fall 2017 and summer 2018. For that we thank Kelly Hogan and Interim Chancellor Kevin Guskiewicz for encouraging Bruno and Steinwand to develop the Seafood Forensics class. The project was also partially funded by the National Science Foundation (OCE #1737071 to JB). We thank Christopher Martin, Chris Willett, and Sabrina Burmeister for sharing their lab space and equipment with us. We also thank the many students who took the class, and collected and processed samples. For spring 2017, we acknowledge Moza Hamud, Meredith McNairy, and Rachel Peterson. For summer 2017, we recognize Aravindhkrishna Ajithkumar, Alanna Dai, Jonathan Dolan, Justin Freeman, Saidou Jallow, Hyun Kim, Matthew Logan, Assem Patel, Grace Tan, and Zachary Young. For fall 2017, we recognize Hanan Alazzam, Alexandra Barry, Christina D’Ovidio, Daniel Efird, Aaron Friedman, Tate Giddens, Mike Grossi, Amaya Martinez, Baily Mcinnes, Kirsi Oldenburg, Steve Park, Farhin Shaikh, Grace Steinman, Nicolas Tobar, Bren Woods. For summer 2018, we recognize Hannah Austin, Jasmine Barnes, Kane Cooper, Julianna Evans, Morgan Korzik, Cassidy Manzonelli, Megan Ochs, and Carlos Urquilla.

## References

Avila J. Antibiotics Illegal in the US Found in Samples of Foreign Shrimp. ABC News Network. 2012 [cited 15 March 2019]. Available from: https://abcnews.go.com/Health/antibiotics-illegal-us-found-samples-foreign-shrimp/story?id=16344514#.UH8T9Rj_Slc

Cawthorn DM, Baillie C, Mariani S. Generic names and mislabeling conceal high species diversity in global fisheries markets. Conservation Letters. 2018; e12573. doi.org/10.1111/conl.12573

Clucas I. A study of the options for utilization of bycatch and discards form marine capture fisheries. FAO Fisheries Circular. No 928. Rome, FAO. 1997; 59p. Available from: http://www.fao.org/3/W6602E/w6602E00.htm

Cox CE, Jones CD, Wares JP, Castillo KD, Mcfield MD, Bruno JF. Genetic testing reveals some mislabeling but general compliance with a ban on herbivorous fish harvesting in Belize. Conservation Letters. 2013; 6(2), 132–140. doi.org/10.1111/j.1755-263X.2012.00286.x

Fofonoff PW, Ruiz GM, Steves B, Simkanin C, Carlton JT. Penaeus vannamei. National Exotic Marine and Estuarine Species Information System. 2018 [cited 15 March 2019]. Available from: https://invasions.si.edu/nemesis/browseDB/SpeciesSummary.jsp?TSN=551682

Garrity-Blake B, Nash B. An inventory of North Carolina fish houses: A five-year update. North Carolina Sea Grant. 2012 [cited 15 March 2019]. Available from: www.ncseagrant.org/s/fhouse-2012

Holmstrom K, Graslund S, Wahlstrom A, Poungshompoo S, Bengtsson BE, Kautsky N. Antibiotic use in shrimp farming and implications for environmental impacts and human health. International Journal of Food Science and Technology. 2003; 38, 255–266. doi.org/10.1046/j.1365-2621.2003.00671.x

Kittinger JN, Teh LCL, Allison EH, Bennett NJ, Crowder LB, Finkbeiner, EM, et. al. Committing to socially responsible seafood: Science. 2017; 356(6341), 912–913. doi.org/10.1126/science.aam9969

Kros JF, Rowe WJ. A Supply Chain Analysis of North Carolina’s Commercial Fishing Industry. North Carolina Rural Economic Development Center. 2013 [cited 15 March 2019]. Available from: https://ncseagrant.ncsu.edu/ncseagrant_docs/products/2010s/supply_chain_analysis_nc_commercial_fishing.pdf

Lobo AS, Balmford A, Arthur R, Manica A. Commercializing bycatch can push a fishery beyond economic extinction. Conservation Letters. 2010; 3: 277–285. doi:10.1111/j.1755-263X.2010.00117.x

Marko PB, Lee SC, Rice AM, Gramling JM, Fitzhenry TM, McAlister JS, et al. Mislabeling of a depleted reef fish. Nature Publishing Group. 2004; 430. doi.org/10.1038/430309b

Marko PB, Nance HA, Van Den Hurk P. Seafood substitutions obscure patterns of mercury contamination in patagonian toothfish (dissostichus eleginoides) or “Chilean Sea Bass.” PLoS ONE. 2014; 9(8), 6–10. doi.org/10.1371/journal.pone.0104140

Marschke M, Vandergeest P. Slavery scandals: Unpacking labour challenges and policy responses within the off-shore fisheries sector. Mar. Policy. 2016: 68, 39–46. doi.org/10.1016/j.marpol.2016.02.009

NOAA. White Shrimp. National Oceanic and Atmospheric Administration. 2018 [cited 15 March 2019]. Available from: https://www.fisheries.noaa.gov/species/white-shrimp

NC Catch. 2006-2018 [cited 15 March 2019]. NC Catch. Available from: https://nccatch.org/pages/our-mission

N.C. Division of Marine Fisheries. N.C. Shrimp. North Carolina Department of Environmental Quality. Raleigh, NC. [cited 15 March 2019]. Available from: http://portal.ncdenr.org/web/mf/nc-shrimp

N.C. Division of Marine Fisheries. 2017 Annual Fish Dealer Report. North Carolina Department of Environmental Quality. Raleigh, NC. 2018 [cited 15 March 2019]. Available from: http://portal.ncdenr.org/c/document_library/get_file?p_l_id=1169848&folderId=30473624&name=DLFE-138254.pdf

National Marine Fisheries Service (NMFS). Fisheries of the United States, 2014. U.S.Department of Commerce, NOAA Current Fishery Statistics No.2014. 2015 [cited 15 March 2019]. Available from: https://www.st.nmfs.noaa.gov/commercial-fisheries/fus/fus14/index

National Marine Fisheries Service (NMFS). Cumulative Trade Data by Product. Fisheries Statistics Division, NOAA Fisheries. [cited 15 March 2019] Available from: https://www.st.nmfs.noaa.gov/commercial-fisheries/foreign-trade/applications/trade-by-product

Paul B, Vogl CR. Impacts of shrimp farming in Bangladesh: Challenges and alternatives. Ocean and Coastal Management. 2011; 54(3), 201–211. doi.org/10.1016/j.ocecoaman.2010.12.001

Staffen CF, Staffen MD, Becker ML, Löfgren SE, Muniz Y, de Freitas R, Marrero AR. DNA barcoding reveals the mislabeling of fish in a popular tourist destination in Brazil. PeerJ. 2017; 5, e4006. doi:10.7717/peerj.4006

Spencer ET and Bruno JF. Fishy Business: Red snapper mislabeling in the southeastern United States. In review.

The Associated Press. Shrimp rise as overall North Carolina commercial catch dips. The Seattle Times. 2018 [cited 15 March 2019]. Available from: https://www.seattletimes.com/nation-world/apxshrimp-rise-as-overall-north-carolina-commercial-catch-dips/

Warner AK, Ph D, Golden R, Lowell B, Disla C, Savitz J. Shrimp: Oceana Reveals Misrepresentation of America’s Favorite Seafood, 2014; 1–46. [cited 15 March 2019]. Available from: https://oceana.org/news-media/publications/reports/shrimpfraud

Warner K, Lowell B, Geren S, Talmage S. Deceptive Dishes□: Seafood Swaps Found Worldwide. Oceana. 2016; 1–21. [cited 15 March 2019]. Available from: https://usa.oceana.org/sites/default/files/global_fraud_report_final_low-res.pdf

Willette DA, Simmonds SE, Cheng SH, Esteves S, Kane TL, Nuetzel H, et al. Using DNA barcoding to track seafood mislabeling in Los Angeles restaurants. Conservation Biology. 2017; 31(5), 1076–1085. doi.org/10.1111/cobi.12888

WoRMS Editorial Board. Litopenaeus setiferus (Linnaeus, 1767). World Register of Marine Species. 2019 [cited 15 March 2019]. Available from: http://www.marinespecies.org/aphia.php?p=taxdetails&id=158336#sources

